# SwimmingIndividuals: A High-Performance Agent-Based Model for Marine Ecosystems

**DOI:** 10.1101/2025.10.13.681996

**Authors:** Matthew S. Woodstock, Jann Paul Mattern, Zhen Wu, Gregory L. Britten

## Abstract

Mechanistic models are essential for projecting ecosystem responses to novel conditions, yet the application and biological realism of agent-based models (ABMs) has often been limited by computational constraints. This paper introduces *SwimmingIndividuals*, an open-source agent-based model (ABM) framework that advances our ability to simulate complex biological processes at large ecological scales. The software’s main contribution is its suite of mechanistic sub-models that govern the full life cycle of each agent, including physics-based visual predation, adaptive behaviors such as diel vertical migration, and a flexible bioenergetics engine with multiple taxa-specific equations. These detailed biological processes are made computationally tractable for large populations through a hybrid CPU/GPU architecture written in the Julia language. We demonstrate the framework’s capabilities through three targeted simulations. The model successfully reproduced complex ecological phenomena as emergent properties, including diel vertical migration, cetacean diving patterns, size-based trophic dynamics, population-level growth curves, and stock-recruitment relationships. Furthermore, long-term simulations generated realistic population dynamics and quantified the impacts of different fishery harvest control rules. By coupling high-resolution biological realism with technical scalability, *SwimmingIndividuals* provides a powerful and flexible tool for a wide range of ecological inquiries. It can be used to conduct in-silico experiments into the ecosystem-scale impacts of environmental disturbances, test fundamental ecological theory, and evaluate the efficacy of complex, ecosystem-based management strategies. This framework advances our ability to build a more mechanistic understanding of marine ecosystem resilience in a changing world.

**Author Summary:** To understand and predict how marine ecosystems function, we need to account for the decisions and life histories of the individual animals within them. We have developed a new open-source software tool, *SwimmingIndividuals*, that acts as a “virtual laboratory” for marine ecology and fisheries science. This software allows us to create large, realistic simulations with millions of virtual organisms, from multiple taxa and species, each with its own unique set of biological traits. These modeled animals make decisions based on their internal state (e.g., hunger) and their perception of the surrounding environment, such as light and temperature. By simulating these individual actions at scale, our software allows us to see emergent, complex, ecosystem-level patterns, like population dynamics and food web structure. *SwimmingIndividuals* provides the scientific community with a powerful tool to investigate fundamental biological questions, explore “what-if” scenarios for fisheries management, and forecast how marine life might respond to future environmental changes.

## Introduction

Predicting the dynamics of marine ecosystems is a critical challenge, as the cumulative effect of individual-scale processes produces emergent, often non-linear dynamics at population and community scales [1]. Capturing this complexity is essential for conservation and management, particularly under the unprecedented pressures from climate change and fishing [2]. Agent-based models (ABMs) represent a fundamentally different approach, modeling individuals as discrete, autonomous entities with unique state variables and behavioral rules [3]. This mechanistic approach is powerful because population-level responses, such as shifts in distribution or recruitment, are not assumed but instead arise directly from individuals responding to their local environment based on physiological and behavioral principles [4].

Traditional ecological models often rely on fitting statistical relationships to historical data. This approach is inherently limited in its ability to provide mechanistic explanations for population changes and becomes highly sensitive to misspecification when used to extrapolate into novel, non-analog conditions created by environmental changes [5, 6]. Although ABMs are better suited for such projections, they have historically faced significant computational limitations that restrict them to smaller spatial scales and fewer agents [7]. These constraints often force the use of simplified rules for key processes that can limit biological realism; for example, an agent’s perception may be a fixed radius rather than being mechanistically determined by light and physiology [8], or its bioenergetics may not be flexible enough to capture diverse metabolic and behavioral strategies [9].

To address these limitations, we developed *SwimmingIndividuals*, a mechanistic model that leverages modern high-performance computing architectures to increase biological realism. Our framework advances previous life-cycle ABMs by bridging the gap between detailed individual processes and large-scale ecosystem simulations. The major advance is the coupling of a scalable hybrid CPU/GPU architecture with a suite of mechanistic sub-models, including a flexible Wisconsin-style bioenergetics model [10] and a predation sequence based on physics-based visual detection. This allows the model to simulate entire life histories governed by complex rules at scales relevant to both theoretical questions and fisheries management. In this paper, we describe the structure of *SwimmingIndividuals* and showcase its ability to reproduce complex ecological phenomena through a series of worked examples, providing a flexible, open-source tool for both testing fundamental ecological theory and exploring applied, conservation-based management strategies.

### Design and Implementation

#### Overview and Design Concepts

*SwimmingIndividuals* simulates the behavior, interactions, movement, and bioenergetics of multiple marine species within a dynamic, spatially explicit environment. The model is designed to explore how individual-scale physiological and behavioral responses to environmental conditions, predator-prey fields, and fishing pressure scale up to produce emergent population- and ecosystem-scale patterns. The model consists of four primary entities: focal species agents, gridded resource patches, a dynamic abiotic environment, and fisheries (Fig 1).

**Fig 1.**
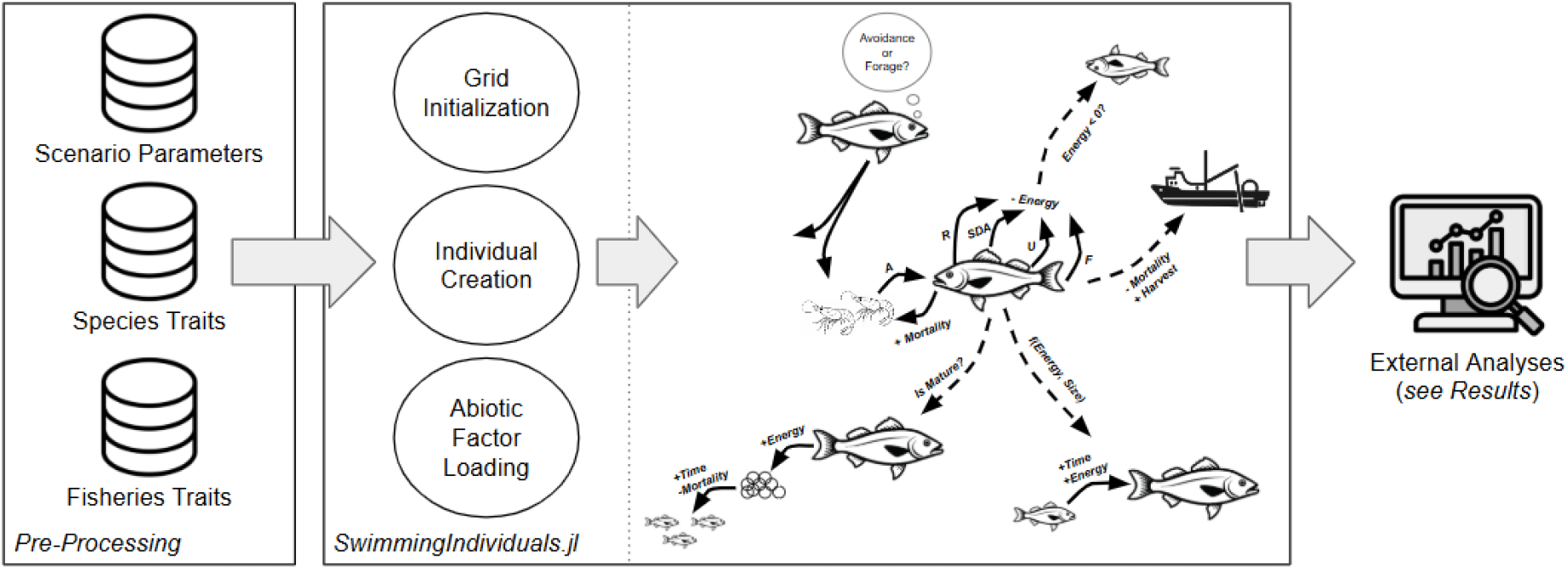
A schematic of the *SwimmingIndividuals* model that represents the agent configuration (traits) required, the initial steps that create the simulated ecosystem, and the behavioral, bioenergetic, and fisheries processes that govern the ABMs and produce results. In this schematic, the fish represents an agent, while the shrimp could either be represented as an agent or resource patch. Each arrow points in the direction of energy transfer. A = assimilated energy, R = respired energy, SDA = specific dynamic action, U = egested energy, F = excreted energy.

Population-level patterns are an emergent property of the cumulative actions of all agents responding to their local conditions. Agents in a *SwimmingIndividuals* model represent a collection of like-size individuals, akin to “superindividuals” in other agent-based models. An agent’s perception of its environment is modeled mechanistically, with its visual range calculated dynamically based on its size, depth, and ambient light. Behavior is adaptive; for example, the time an agent spends foraging is regulated by its internal state (i.e., gut fullness) and the local availability of prey. The implicit objective of each agent is to maximize fitness by surviving and accumulating surplus energy for growth and reproduction. Stochasticity is incorporated into key processes, including habitat selection and prey capture, to reflect the randomness inherent in natural systems.

#### Simulation Modules

The simulation proceeds in discrete time steps, governed by a series of modules that update the state of each agent and the environment (Fig 2).

**Fig 2.**
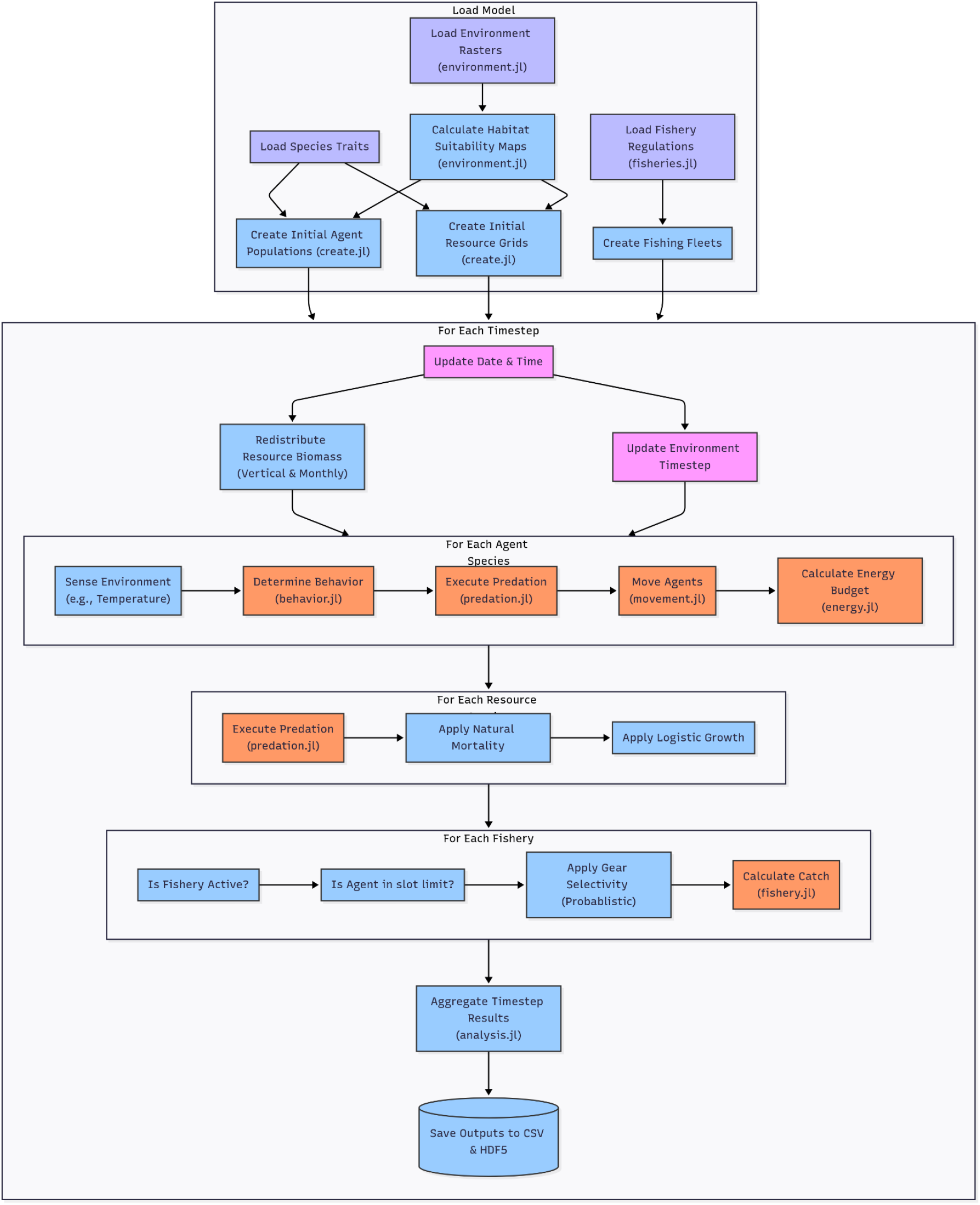
Workflow of a *SwimmingIndividuals* model showing data inputs (purple), submodels (orange), necessary processes and checks (blue), and ecosystem state changes (pink).

### Initialization

At the start of a simulation, the model creates focal species agents, resource patches, the gridded environment, and fishing fleets. Agents and resources are placed in the model domain based on habitat capacity maps, and their initial properties (e.g., length, depth, location) are drawn from user-defined statistical distributions.

### Movement

Agent movement is determined by its behavioral state. The default behavior is to move towards higher-quality habitat, with a path calculated by a pathfinding algorithm. This habitat-seeking movement can be overridden by specialized vertical movements that simulate behaviors such as diel vertical migration (DVM) or the diving patterns of marine mammals.

### Predation

The predation submodel manages interactions through a sequence of mechanistic steps. A predator first identifies potential prey within its dynamically calculated visual range and appropriate predator-prey size ratio. It then performs a localized search to select the best target (currently the nearest potential prey without species’ preference) and attempts consumption. Success is limited by a species-specific handling time and the predator’s current gut fullness, creating an emergent functional response.

### Bioenergetics

The bioenergetics submodel calculates a full energy budget for each agent based on a flexible, Wisconsin-style framework [10]. It accounts for energy gained from consumption and costs from respiration, excretion, and activity. The model includes multiple taxa-specific respiration equations to accommodate diverse metabolic strategies (e.g., fishes, cetaceans, deep-sea). Surplus energy is allocated to somatic growth and, for mature individuals, reproduction. Negative energy balance over time can lead to starvation mortality.

### Fisheries

The fisheries submodel applies fishing mortality to target and bycatch species. Biomass is removed according to fleet-specific rules, including quotas, seasons, bag limits, and gear-based size selectivity. The model tracks catch, effort, and fishing mortality over space and time.

## Results

To demonstrate the capabilities of the *SwimmingIndividuals* framework, we conducted three targeted simulations in the Gulf of Mexico: a one-week “DVM” scenario with two vertically migrating species; a two-day “Diver” scenario with two deep-diving cetacean analogues; and a 50-year “Mackerel” scenario with King Mackerel (*Scomberomorus cavalla*) and nineteen resource groups. Below, we showcase the model’s ability to reproduce complex ecological phenomena as emergent properties using results from the three scenarios.

### Emergent Movement Patterns

The model’s ability to simulate complex, state-driven movement is a core feature. Simulations from the DVM scenario reproduced the daily vertical migrations of both strong and weak migrant species, which descend during the day and ascend at night to forage (Fig 3A). The model also generated the repeated, deep foraging dives and subsequent surface intervals characteristic of cetaceans (Fig 3B), allowing for outputs like daily travel distance (Fig 3C).

**Fig 3.**
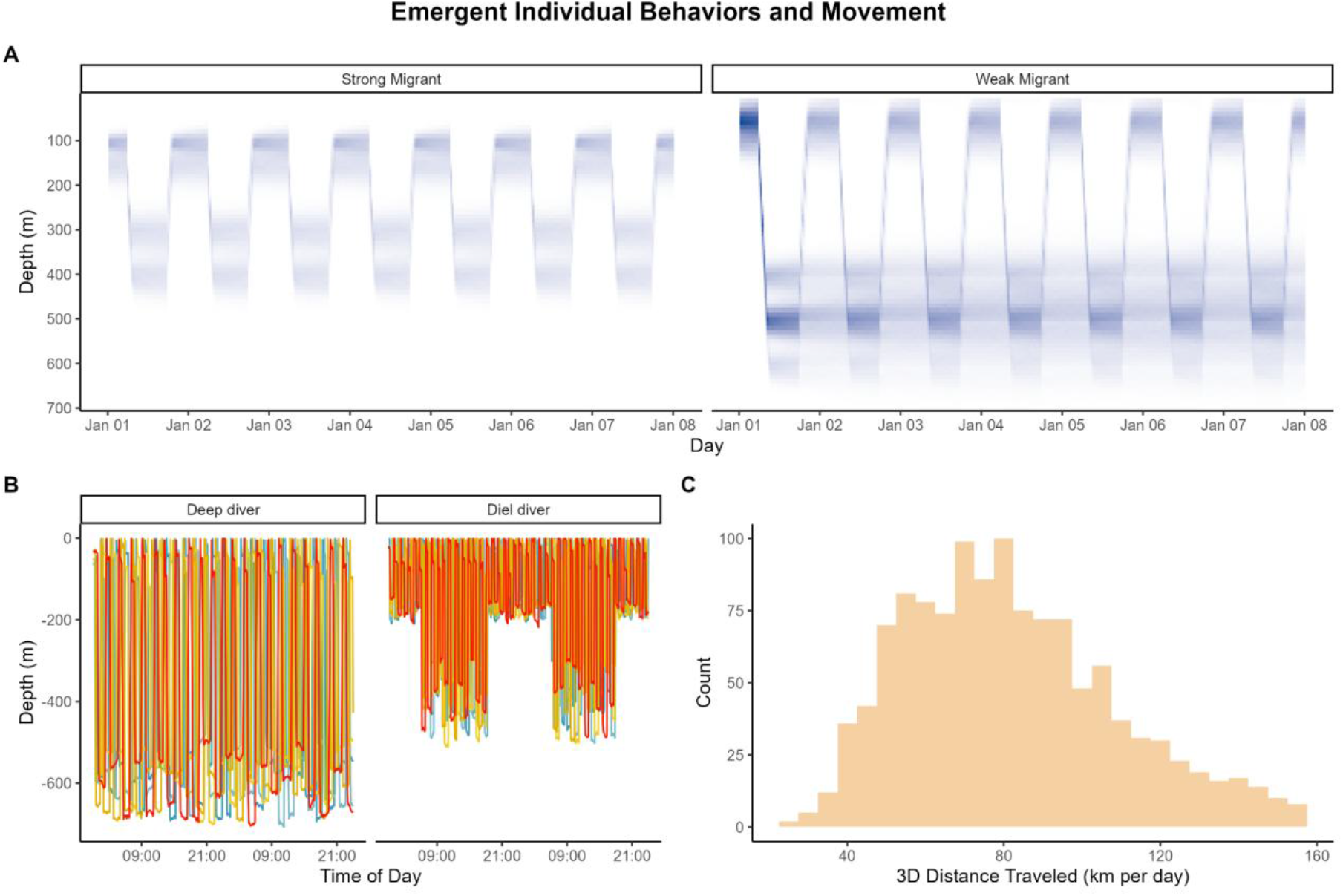
A representation of the emergent movement patterns of A) strong and weak diel vertical migrant populations, B) two species of deep-diving cetaceans (5 individuals only); one that is a deep diver (e.g., sperm whale) and another that has a diel dive pattern (e.g., Rice’s whale), and C) a histogram of the 3D movement distance of each agent for the deep-diving cetacean. Colors in panel A correspond to a gradient of dense aggregations of individuals (dark blue) and unoccupied depths (white).

Large-scale patterns, such as seasonal migrations, emerge organically from localized behavioral rules without being explicitly programmed. This is achieved through a habitat-seeking behavior where each agent assesses its immediate surroundings and makes incremental movements towards location with higher habitat quality. These habitat quality scores are dynamic, shifting throughout the year as the environmental data (e.g., temperature) changes monthly. While no single agent is aware of a migratory path, the cumulative effect of each agent independently making localized, optimal decisions results in a cohesive and large-scale shift in the population’s spatial distribution (Fig 4).

**Fig 4.**
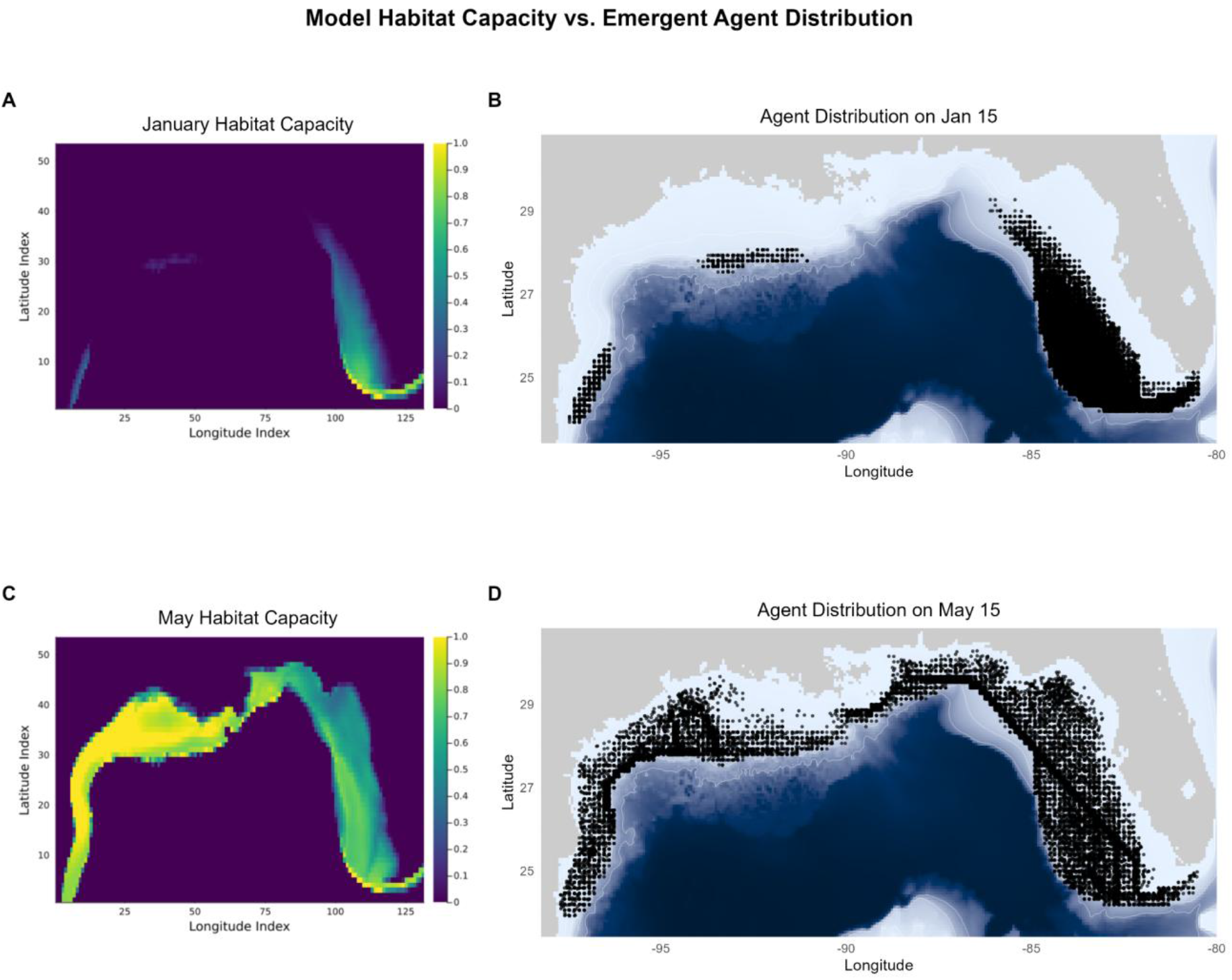
The matchup between habitat capacity (A and C) and agent distributions (B and D) in different months of the year, as determined by incremental movements by each agent. Panels A and C are represented on the model grid, whereas panels B and D reflect individuals at particular longitude and latitude coordinates. *SwimmingIndividuals* tracks both grid locations and geographic coordinates for habitat seeking, movement, prey searching, and bioenergetic calculation functionalities.

### Predator-Prey and Trophic Dynamics

The multi-species Mackerel scenario allows for a comprehensive analysis of trophic linkages. Model outputs can be used to construct aggregated food webs (Fig 5A) and reveal size-based shifts in diet (Fig 5B). Further, spatially explicit consumption matrices can identify predation “hotspots” where predator-prey interactions are concentrated due to overlapping habitat preferences (Fig 5C).

**Fig 5.**
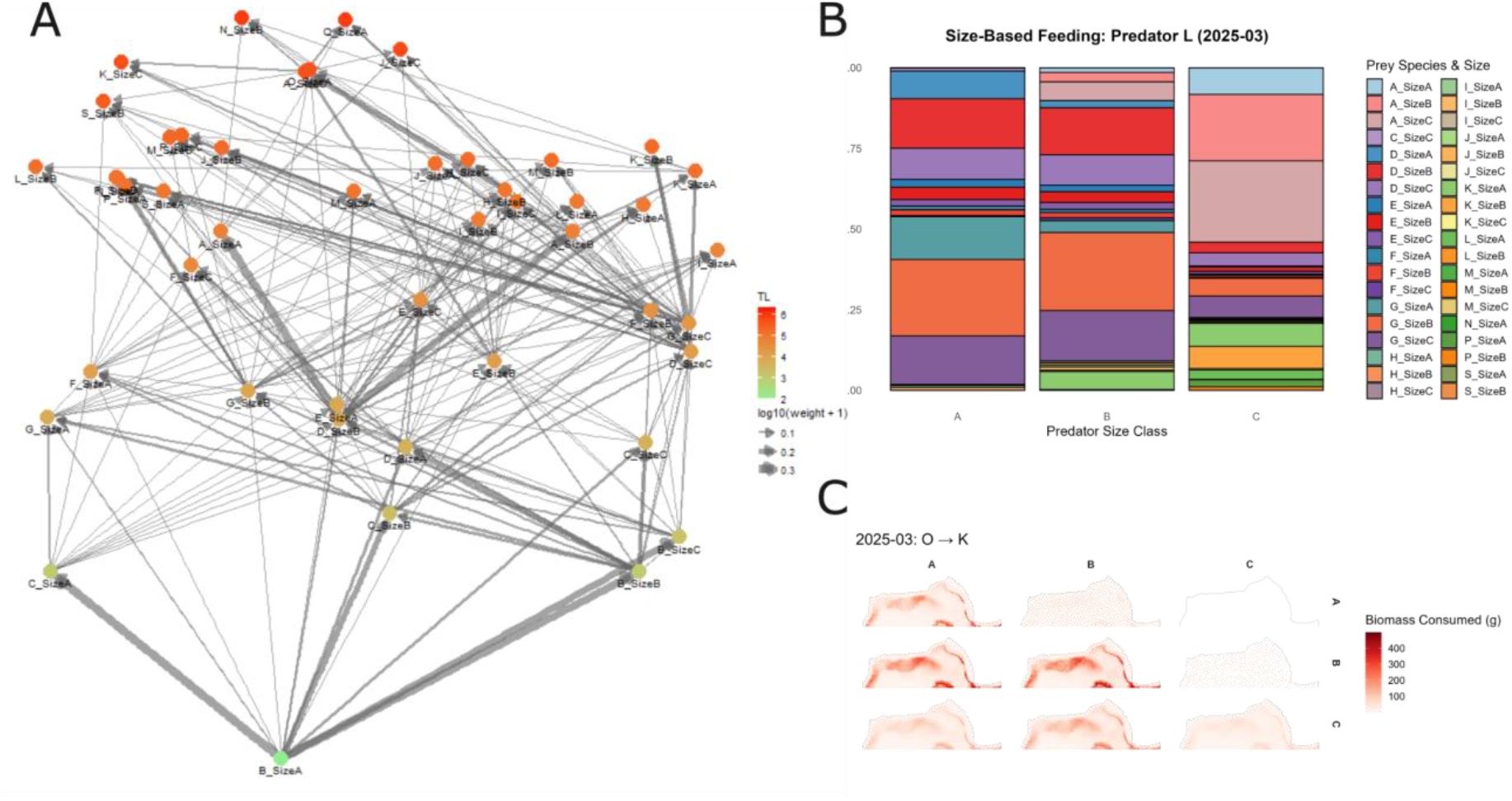
Representations of predator-prey dynamics that emerge from a simulation with one focal species (species A) and 19 resource species simulation. All plots were created from daily outputs that were aggregated to a monthly scale for brevity. In this example, panel A) The diet matrix can construct timestep and grid-cell specific food webs that can be aggregated post-hoc to the appropriate spatial and temporal scale. Fractional trophic levels (y-axis) were calculated post-hoc assuming size classes that ate nothing were primary consumers (i.e., trophic level = 2). The line weighting corresponds to the magnitude of energy transfer. B) Predator L (e.g., dolphinfish) has size-based differences in prey composition that correspond to predator-prey encounters and predator-prey size relationships. Size classes A-C correspond to small to large individuals of the same species. C) Spatially explicit biomass consumption from a predator species, O (e.g., sailfish; columns), to a prey species, K (e.g., little tunny; rows), delineated by A (small) to C (large) size classes.

### Life History and Bioenergetics

The bioenergetics submodel drives individual growth, reproduction, and feeding rates, producing a wide range of life history outputs. These include fine-scale individual metrics like relative energy reserves (Fig 6A), metabolic costs (Fig 6B), and gut fullness cycles (Fig 6C). At the population level, the model generates emergent von Bertalanffy-style growth curves (Fig 6D) and stock-recruitment relationships (Fig 6E), which can be validated against empirical data.

**Fig 6.**
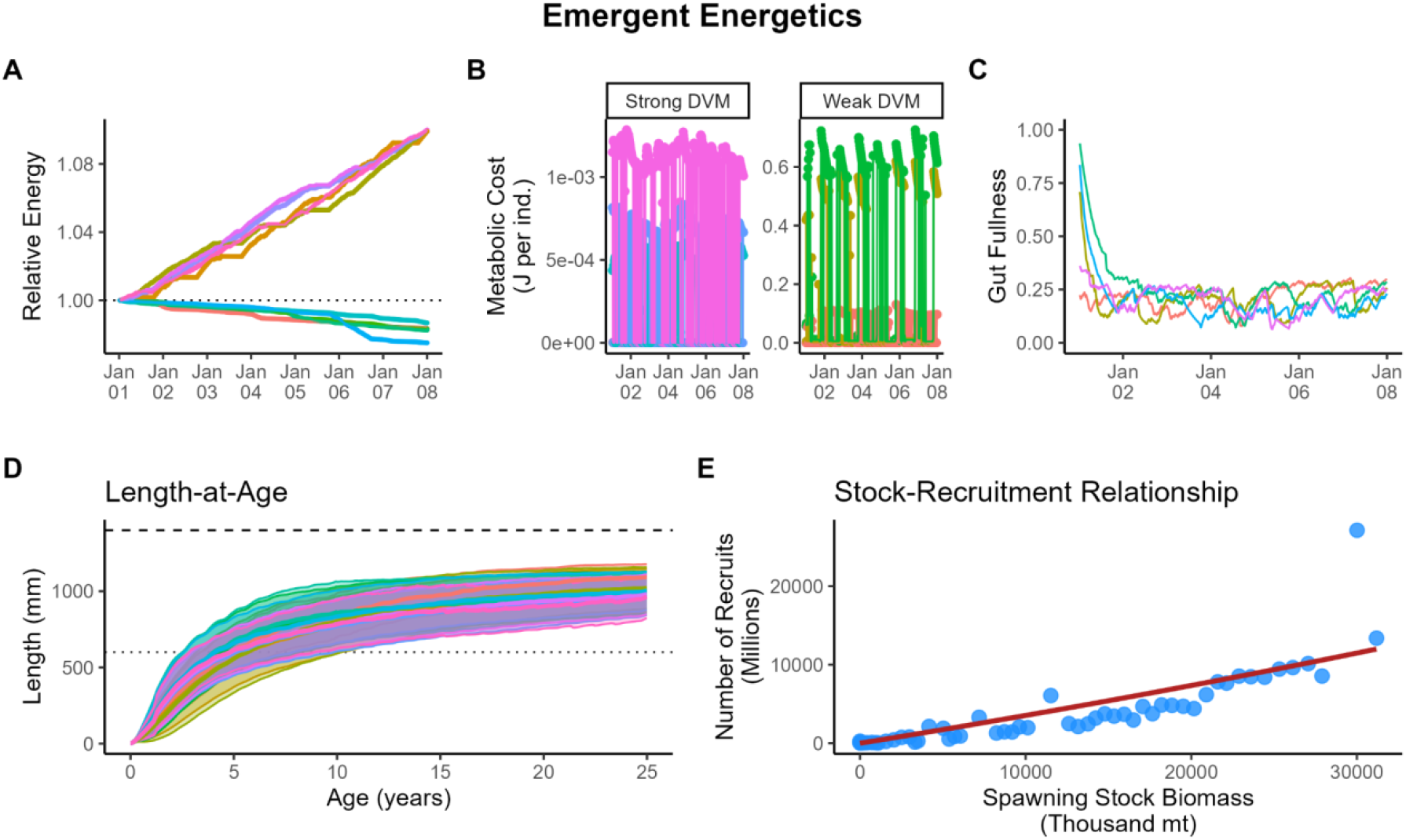
Examples of energetics-based analyses that can be derived from a *SwimmingIndividuals* simulation. A) Relative trends in energy reserve storage of 10 select individuals (colors) from the diel vertical migration model run. B) Metabolic costs (J per timestep) over time in the diel vertical migration model run for 3 select individuals (colors) each facet. Y-axes are different because of differences in body size between the two species. C) Gut fullness trends among five strong migrant individuals (colors) in the diel vertical migration simulation. Reduced gut fullness over time promotes a higher likelihood of feeding, assuming prey are available. D) Length-at-age curve among distinct generations (colors) in the mackerel simulation. Solid middle lines are the mean length-at-age (mm-scale), surrounded by ± 1 standard deviation. The dotted line (600 mm) is the length at maturity and the dashed line (1400 mm) is the length at infinity. This plot was cut off at 25 years of age and certain applications may call for an age-based senescence, rather than the current length-based senescence. E) The stock-recruitment relationship of mackerel; annual recruits (Age-0 individuals) to annual spawning stock biomass (sum of biomass of mature individuals). Each point represents a different year and the model results were fit to a Beverton-Holt curve.

### Population Dynamics

The culmination of individual life histories gives rise to realistic population-level patterns. In the Mackerel scenario, total abundance fluctuated with seasonal spawning cycles (Fig 7A), leading to shifts in the population’s size-frequency distribution over the year (Fig 7B). Consequently, the mean size of an individual in the population changed over time, often inversely related to recruitment pulses (Fig 7C). The biomass of resource species also fluctuated based on both user-defined seasonal growth and internal food-web dynamics (Fig 7D).

**Fig 7.**
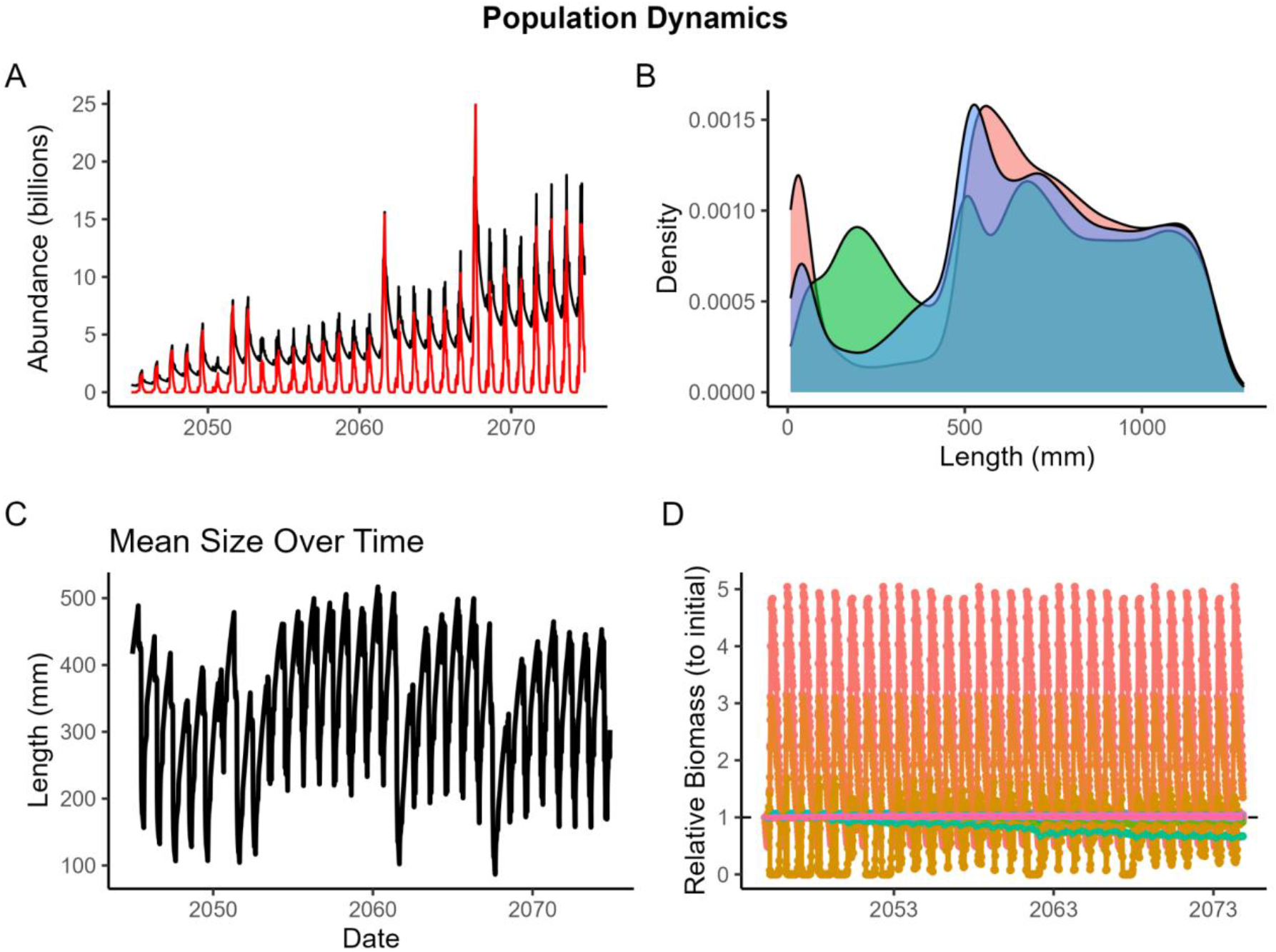
Several population dynamics metrics from a *SwimmingIndividuals* model, including; A) total abundance (black) and larval abundance (red) emergent from a seasonal spawning period and low predator density; B) size-frequency distributions of a focal species population at three slices in time; C) the mean size of all individuals in the population that is inversely related to peak spawning periods; and D) the relative biomass trends of each resource species that fluctuates seasonally with seasonal growth patterns and predation mortality.

### Fisheries Impacts

*SwimmingIndividuals* can serve as a testbed for management strategies by simulating the impacts of fishing. In the Mackerel scenario, distinct harvest control rules for five different fisheries resulted in different rates of quota attainment (Fig 8A) and variable mean lengths of caught fish (Fig 8B). The model tracks the resulting time-varying fishing mortality for specific size classes (Fig 8C) and can map its spatial distribution, providing a tool to evaluate the efficacy of time-area closures (Fig 8D).

**Fig 8.**
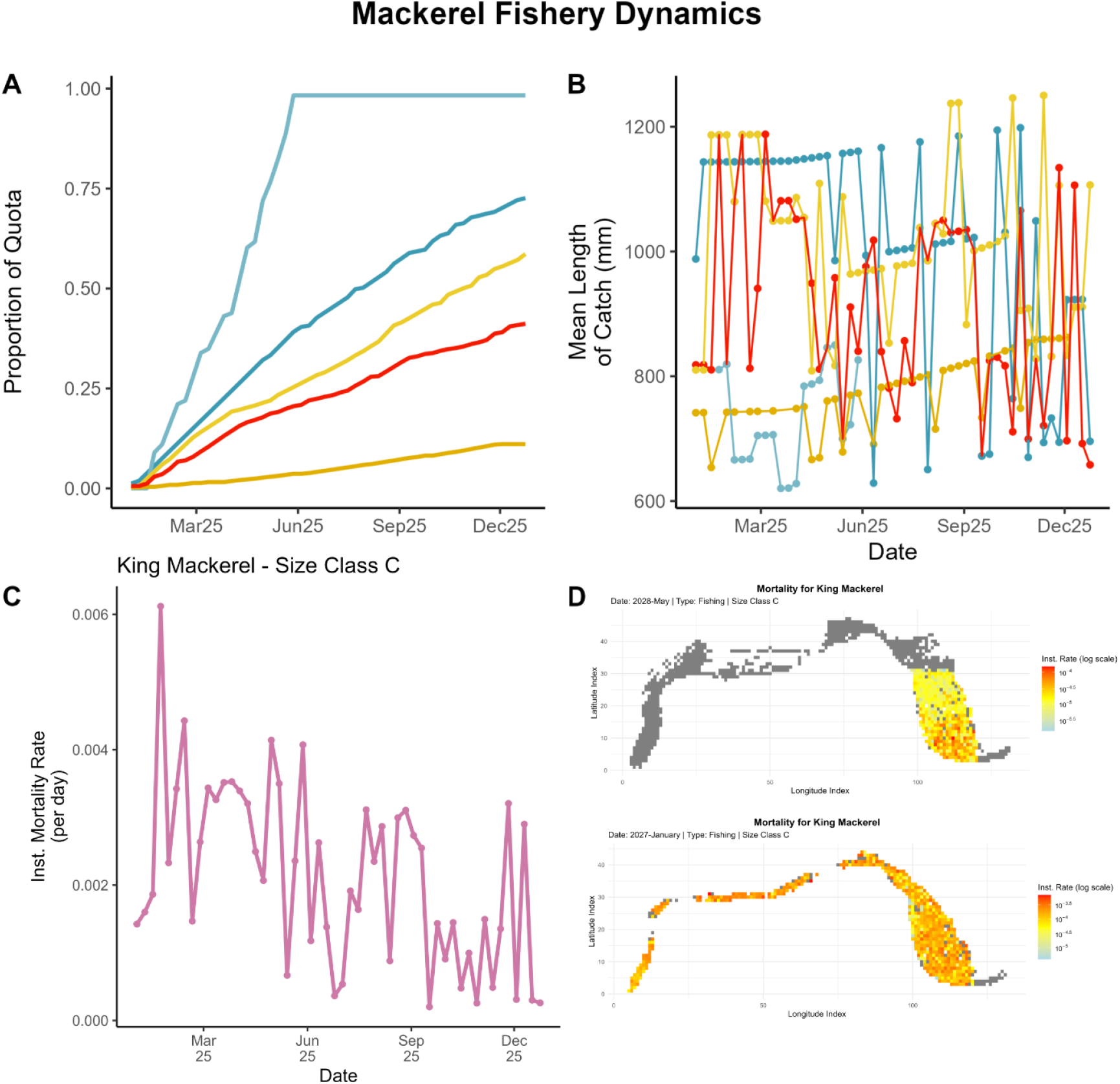
Fisheries outputs from a *SwimmingIndividuals* simulation showing A) the proportion of quota usage throughout the first year of the simulation for each fishing fleet (colors). Different fishing fleets had different fishing areas, quotas, and selectivity. B) The mean length of the catch for each fishing fleet (same colors as A), C) time-varying fishing mortality for the largest size class of a focal species in the first year of the simulation, and D) the spatial distribution of fishing mortality for two distinct months for the largest size class. Fishing mortality values are log-transformed instantaneous rates per grid cell. Only grid cells occupied by King Mackerel are displayed, but the plot generally outlines the coastal region (0–400m depth).

## Availability and Future Directions

The *SwimmingIndividuals* framework contributes to the development of mechanistic tools for marine ecosystem modeling. By coupling a detailed, agent-based representation of organismal life history with a modern computing architecture, the model provides a powerful platform for in-silico experimentation [11, 5]. This process-based approach is vital for projecting the impacts of climate change, where novel environmental conditions can render historical correlations unreliable [6, 12]. The model’s applications span both applied conservation and theoretical ecology. For conservation, its most direct uses are to identify biological parameters for assessments (e.g., age-length curves, stock-recruitment relationships) and as a Management Strategy Evaluation (MSE) tool [13], allowing for the testing of complex regulations to predict cascading effects on population structure and the broader food web [14, 15]. For theoretical ecology, the framework serves as a tool for investigating how individual heterogeneity scales up to affect population stability [16], or testing hypotheses about the mechanisms driving spatial patterns in predator-prey interactions.

The full, open-source code for *SwimmingIndividuals* is publicly available on Github under an MIT license with a user manual. Potential users can copy the source code, make issue statements for the development team, and suggest additions to the existing model structure for future releases. The repository includes all source code, input data files for the case studies, and R scripts for post-processing and analysis, which can be supplemented by the community. The software can be run on any operating system that supports Julia and installs the necessary dependencies listed in the *model*.*jl* source code file (all Julia packages).

The current limitations of *SwimmingIndividuals* v1.0 highlight several avenues for future development and community contribution. For instance, the model does not track larval dynamics to limit the computational expense of a model run, although individual larvae can be consumed by predators. The absence of passive larval dispersal presents an opportunity to integrate the model with hydrodynamic simulations to investigate population connectivity [17]. The lack of complex internal schooling dynamics invites extensions to explore the emergent anti-predator and foraging benefits of group behavior [18]. Furthermore, the model’s fleet-wide, static harvest rules could be expanded to have a more comprehensive, agent-based vessel structure to allow for within-fleet differences in fisher behavior and harvest. Addressing these gaps through collaborative, open-source development will be essential for advancing mechanistic ecosystem modeling.

## Acknowledgments

We are grateful to Alex Norelli for commenting on an early version of this manuscript. MSW was partially supported by the Woods Hole Oceanographic Institution Postdoctoral Scholar Award. This research was carried out in part under the auspices of the Cooperative Institute for Marine and Atmospheric Studies (CIMAS), a Cooperative Institute of the University of Miami and the National Oceanic and Atmospheric Administration, cooperative agreement #NA20OAR4320472.

